# Development of Visual Response Selectivity in Cortical GABAergic Interneurons

**DOI:** 10.1101/2021.07.21.453281

**Authors:** Jeremy T. Chang, David Fitzpatrick

**Affiliations:** Department of Functional Architecture and Development of Cerebral Cortex, Max Planck Florida Institute for Neuroscience, Jupiter, Florida, USA

## Abstract

The visual cortex of carnivores and primates displays a high degree of modular network organization characterized by local clustering and structured long-range correlations of activity and functional properties. Excitatory networks display modular organization before the onset of sensory experience, but the developmental timeline for modular networks of GABAergic interneurons, remains under-explored. Using *in vivo* calcium imaging of the ferret visual cortex, we find evidence that before visual experience, interneurons display weak orientation tuning and widespread non-specific activation in response to visual stimuli. Modular organization and orientation tuning are evident with as little as one week of visual experience. Furthermore, we find that the development of orientation tuning requires visual experience, while the reduction in widespread network activity does not. Thus, the maturation of inhibitory cortical networks occurs in a delayed, parallel process relative to excitatory neurons.

## Introduction

The onset of visual experience coincides with a period during which cortical circuits are highly plastic. This enables visual experience to drive maturation of the functional properties and network organization of excitatory neurons in the visual cortex, improving orientation and direction selectivity and establishing binocular alignment of response preferences^1–5^. The visual cortex of carnivores and primates has a modular arrangement of excitatory neurons, which is characterized by both the local clustering and long-range patterning of neuronal responses and functional properties such as orientation preference. Modular network organization is evident in excitatory networks before visual experience for both spontaneous activity and visually evoked responses to orientation, suggesting it develops in an experience-independent manner^4,6,7^. Working in concert with these excitatory neurons are inhibitory, GABAergic interneurons (GABA-INs). Recent work in the mature ferret has shown that GABA-INs are orientation selective and tightly coupled to the excitatory network—exhibiting a modular structure, where neighboring inhibitory and excitatory neurons share common orientation preferences^8^. The development of these aligned inhibitory and excitatory networks is a remarkable feat, as excitatory and inhibitory neurons derive from different progenitors and undergo distinct patterns of migration^9^, yet develop well-ordered functional maps. Furthermore, the co-development and interaction of inhibitory and excitatory neurons is an important component of the normal development of cortical function^10,11^, and alterations in inhibitory signaling across development can alter the structure of sensory maps^12^.

Despite the important contribution of GABA-INs in the development of cortical networks, the sequence of events that underlie the emergence of mature interneuron functional properties and the role of visual experience in this process remain poorly understood. Because GABA-INs in the mature mouse visual cortex lack the selective responses and modular structure found in higher mammals^13,14^, it is unclear how the results from developmental studies in mice generalize to other species. Nevertheless, there is evidence for developmental experience-dependent changes in the functional properties of mouse cortical interneurons (becoming more broadly tuned with experience^15,16^), and recent transplantation studies have highlighted the importance of innate, experience-independent mechanisms gating the period for interneuron maturation^16^. Here we examine the contributions of experience-dependent and -independent mechanisms in the maturation of the functional properties and network organization of GABA-INs in the modularly organized ferret visual cortex using two-photon and widefield epifluorescence calcium imaging.

We find that before the onset of visual experience, when excitatory neurons display orientation selectivity^4^, GABA-INs are poorly orientation tuned, and display a high degree of response variability. The immaturity of GABA-IN responses at this time point is also evident at the network-scale where visual stimuli drive widespread patterns of activity lacking the modular structure that dominates mature cortex. These immaturities resolves shortly after the onset of visual experience, and evidence from deprivation experiments suggests that both experience-dependent and independent mechanisms contribute to this process. Thus the development of the functional properties and modular network structure of GABA-INs in visual cortex appears delayed relative to excitatory neurons, a chronology that suggests an instructive role for excitatory networks in the development of inhibitory circuits.

## Results

### GABAergic interneurons develop orientation selectivity after eye opening

Using viral expression of GCaMP6s under the mDlx enhancer, which preferentially labels GABA-INs^8,17^, we assessed single neuron responses in layer 2/3 of ferret visual cortex to the binocular presentation of square drifting gratings with two-photon calcium imaging. We compared 3 groups of animals with different amounts of visual experience: those with no visual experience (at or before the onset of visual experience, designated ‘Naive’); those with a short period of visual experience (between 4 and 7 days of visual experience, designated ‘Brief’; and those with extended visual experience (greater than 8 days after the onset of visual experience, designated ‘Extended’). GABA-INs in the visual cortex of animals with visual experience (both Extended and Brief) exhibited comparably strong orientation selective responses that were not significantly different. (Figure 1D-F, Brief 0.54 ± 0.06 vs Extended 0.52 ± 0.05, p=0.8298, n= 6 vs 5, Mean ± SEM, bootstrap significance test). Modest improvements were observed in the direction selectivity (Figure 1G, Brief 0.16 ± 0.03 vs Extended 0.24 ± 0.03, p=0.0444, n= 6 vs 5, Mean ± SEM, bootstrap significance test). Also, the fraction of significantly orientation tuned cells did not change over this time (Figure 1J, Brief 0.81 ± 0.03 vs Extended 0.64 ± 0.05, p=0.0521, n= 6 vs 5, Mean ± SEM, bootstrap test), and the fraction of cells that were significantly direction tuned only modestly increased over this period (Fraction of cells significantly direction tuned, Brief 0.17 ± 0.02 vs Extended 0.28 ± 0.03, p=0.0214; n= 6 vs 5, Mean ± SEM, bootstrap test). Taken together, these data suggest that the orientation and direction selectivity of GABA-INs is at near mature levels within one week of visual experience.

**Figure 1.**
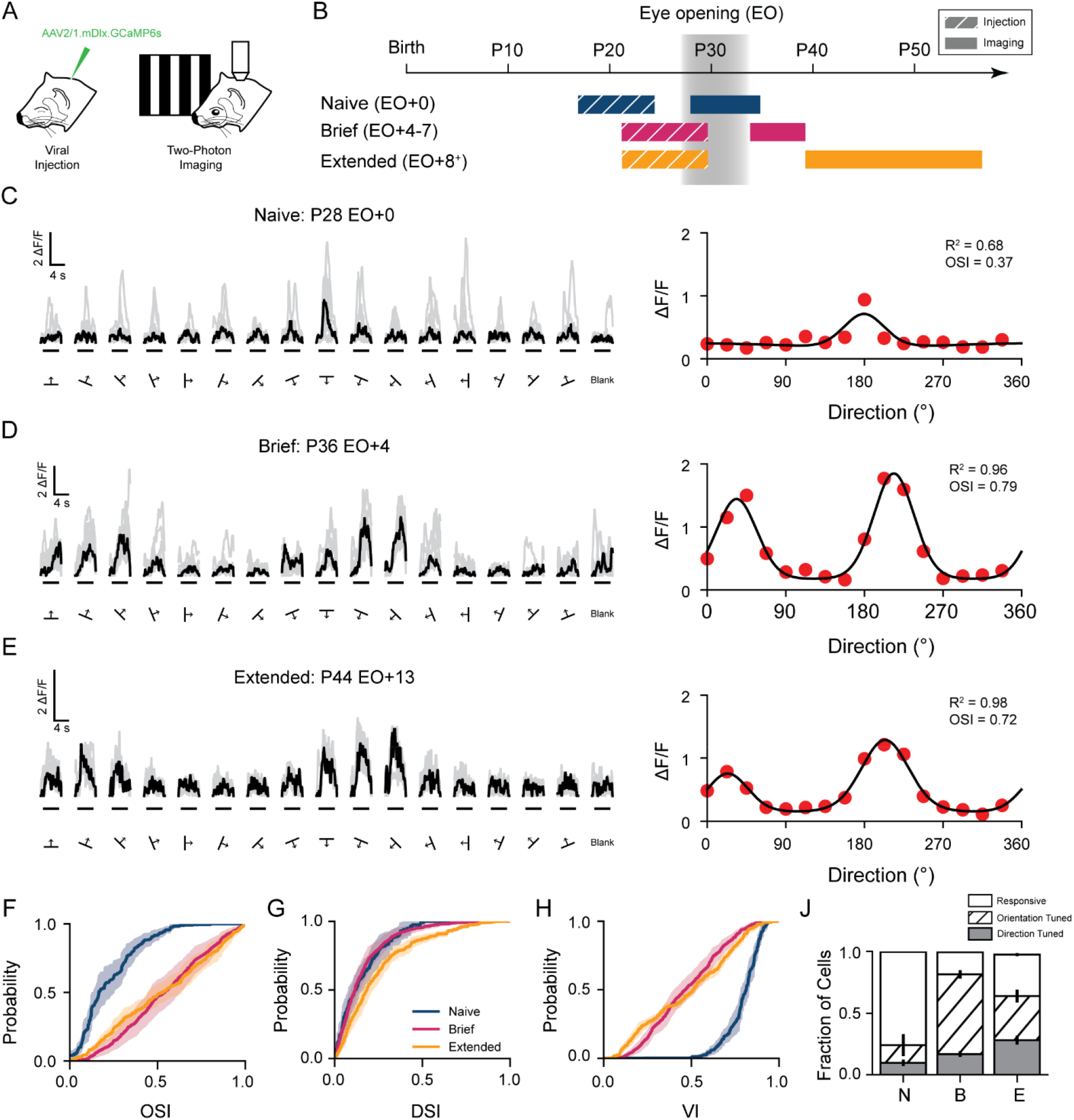
Orientation selectivity develops in interneurons after eye opening. (A) Two-photon imaging of viral expression of GCaMP6s under the mDlx enhancer in ferret visual cortex (B) Three age groups, Naive (EO+0), Brief experience (EO+4-7), and Extended experience (EO+8^+^) (C) Example cellular responses to square drifting gratings (left, black traces denote median response and gray traces denote single trials) and responsive tuning curve (right) for a Naive animal. (D-E) same as (C) but for a Brief and Extended experience animal respectively. (F-H) Cumulative distributions of orientation selectivity index (F), direction selectivity (G), and variability index (H) for Naive (blue, n=7), Brief (magenta, n=6), and Extended (gold, n=5) experience animals. Shaded regions denote SEM. (J) Fraction of cells that are responsive (white), orientation selective (hatched), and direction selective (gray) for Naive (N), Brief (B), and Extended (E) experience animals. Error bars denote SEM.

In contrast, GABA-INs in visually naive animals (Figure 1C, Naive) exhibit only weak orientation selectivity compared to experienced animals, with a mean orientation selectivity that is significantly lower (Figure 1E, Naive 0.22 ± 0.04 vs Brief 0.54 ± 0.06, p =0.0032, n= 7 vs 6; Naive 0.22 ± 0.04 vs Extended 0.52 ± 0.05, p= 0.0028, n= 7 vs 5; Mean ± SEM, bootstrap significance test) The naive group also exhibited a lower percentage of orientation tuned cells (Figure 1J, Naive 0.24 ± 0.09 vs Brief 0.81 ± 0.03, p=0.0344, n= 7 vs 6; Naive 0.24 ± 0.09 vs Extended 0.64 ± 0.05, p= 0.0164, n= 7 vs 5; Mean ± SEM, bootstrap significance test). A small fraction of GABA-INs displayed direction selectivity; however, the overall proportion did not significantly change over the first week of experience (Figure 1G, Naive 0.10 ± 0.03 vs Brief 0.17 ± 0.02, p = 0.0638, n= 7 vs 6, Mean ± SEM, bootstrap significance test), and selectivity only modestly increased over the second week of visual experience (Naive 0.10 ± 0.03 vs Extended 0.28 ± 0.03, p= 0.0040, n= 7 vs 5, Mean ± SEM, bootstrap significance test). The lack of orientation selectivity at eye opening, however, does not reflect a lack of visual responsiveness, as we did not find a significant difference in the percentage of visually responsive cells (Fraction of responsive cells, Naive 0.998 ± 0.001 vs Brief 0.995 ± 0.003, p=0.3412, n= 7 vs 6; Naive 0.998 ± 0.001 vs Extended 0.975 ± 0.013, p=0.0647, n= 7 vs 5; Mean ± SEM, bootstrap test).

One apparent characteristic of visual responses in Naive animals that could contribute to the poor selectivity is the variability in trial-to-trial cellular responses for a stimulus (Figure 1C), a phenomenon that was not seen in animals with experience (Figure 1D, E). To quantify the reliability of cellular responses, we calculated a variability index, a measure of the consistency of cellular tuning curves (see Methods). Naive animals show a higher degree of cellular variability than animals with visual experience (Naive 0.79 ± 0.02 vs Brief 0.48 ± 0.04, p= 0.0004, n= 7 vs 6; Naive 0.79 ± 0.02 vs Extended 0.48 ± 0.02, p= 0.0004, n= 7 vs 5; Mean ± SEM, bootstrap significance test). These findings suggest that before eye opening, GABA-INs are poorly tuned to stimulus orientation in part because of cellular response variability and that rapid changes in the week following the onset of visual experience lead to reductions in variability and mature levels of orientation selectivity.

A property of excitatory neurons that undergoes a dramatic experience-dependent change following eye opening is the alignment of orientation preferences for responses driven by either the contralateral or ipsilateral eye. Before eye opening in ferret, excitatory neurons often have mismatched orientation preferences for each eye that become aligned with visual experience^15^. We thought it would be important to determine whether a mismatch in orientation preferences for the contralateral and ipsilateral eyes contributed to the weak orientation selectivity in the Naive group, since these initial experiments were conducted using binocular stimulation. Using monocular stimulation, we found that almost all GABA-INs in the Naive group responded to the monocular stimulation of the contralateral and ipsilateral eye (Supplementary Figure 1D, H; Fraction of responsive cells, Naive contralateral 0.994 ± 0.003, ipsilateral 0.992 ± 0.03, n= 7, Mean ± SEM). Like the binocular responses, these responses were not well-tuned to orientation or direction (Supplementary Figure 1 A, B, E, F. Fraction of orientation tuned cells Naive contralateral 0.144 ± 0.042, ipsilateral 0.192 ± 0.085; Fraction of direction tuned cells Naive contralateral 0.072 ± 0.03, ipsilateral 0.044 ± 0.016; n=7, Mean ± SEM), supporting our conclusion that GABA-INs in visually naive animals lack orientation selectivity for both monocular and binocular stimulation.

The higher percentage of binocular GABA-INs in the Naive group led us to wonder whether this property was maintained after the onset of experience and whether the emergence of orientation selectivity in GABA-INs exhibited an initial mismatch in orientation selectivity like that found for excitatory neurons that is subsequently refined. We found that GABA-INs appear to maintain a high degree of binocularity across all groups that we examined with no significant shift in monocularity (Supplementary Figure 1I; |ODI| Naive 0.14 ± 0.02 vs Brief 0.16 ± 0.02, p= 0.3216, n= 7 vs 6; Naive 0.14 ± 0.02 vs Extended 0.14 ± 0.01, p= 0.8352, n = 7 vs 5; Mean ± SEM, bootstrap test). Moreover, there was no significant difference in the monocular orientation preference mismatch between the two age groups with visual experience (Supplementary Figure 1L; Ipsilateral vs Contralateral, Brief 8.54 ± 1.17 vs Extended 6.26 ± 1.48, p=0.2116; n=6 vs 5, bootstrap significance), or in the mismatch of monocular and binocular orientation preferences (Supplementary Figure 1J, K; Binocular vs Contralateral, Brief 4.98 ± 0.77 vs Extended 5.15 ± 0.60, p= 0.8418; Binocular vs Ipsilateral, Brief 7.63 ± 1.01 vs Extended 5.13 ± 0.44, p=0.0560; n= 6 vs 5; Mean ± SEM, bootstrap significance).

Taken together, these results indicate that inhibitory and excitatory neurons have fundamentally different developmental progressions for orientation selectivity and binocularly aligned responses. For excitatory neurons, orientation selectivity emerges before experience and visual experience achieves binocular alignment and sharpens tuning. In contrast, orientation selectivity and binocular alignment emerge concurrently in GABA-INs shortly after eye opening.

### Developmental emergence of orientation selective signals in populations of GABAergic interneurons

Although we found that individual GABA-INs in Naive animals are not well-tuned for orientation, we wondered whether there was enough reliability in cellular responses present across the population of interneurons to support a population-level representation of stimulus orientation. To assess this, we measured the trial-to-trial pattern correlation (Figure 2A) of stimulus-evoked responses for the entire population of GABA-INs across all orientations and then compared the values for matched and orthogonal stimuli. In Naive and experienced animals, trial-to-trial correlations for matched stimuli were significantly higher than orthogonal stimuli (Figure 2B, Naive, matched 0.59 ± 0.02 vs orthogonal 0.50 ± 0.04, p=0.0316, n=7; Brief matched 0.71 ± 0.02 vs orthogonal 0.02 ± 0.07, p=0.0006, n= 6; Extended matched 0.76 ± 0.02 vs orthogonal 0.09 ± 0.02, p<0.0001, n=5; Mean ± SEM, Student’s paired t-test). Although the difference in trial-to-trial correlations was significant for Naive animals, the difference in trial-to-trial correlations for matched and orthogonal stimuli was reduced to a fraction of the values found in experienced animals, such that pattern correlations were significantly higher for matched stimuli (Naive 0.59 ± 0.02 vs Brief 0.71 ± 0.02, p= 0.0004, n= 7 vs 6; Naive 0.59 ± 0.02 vs Extended 0.76 ± 0.02, p= 0.0012, n= 7 vs 5; Mean ± SEM, bootstrap test), and significantly lower for orthogonal stimuli (Naive 0.50 ± 0.04 vs Brief 0.03 ± 0.07, p= 0.0016, n= 7 vs 6; Naïve 0.50 ± 0.04 vs Extended 0.09 ± 0.02, p= 0.0010, n= 7 vs 5; Mean ± SEM, bootstrap test). These data demonstrate that oriented stimuli evoke responses in GABA-INs at eye opening, but the pattern of response is only weakly related to stimulus orientation, displaying a higher response similarity across stimuli than animals with visual experience.

**Figure 2.**
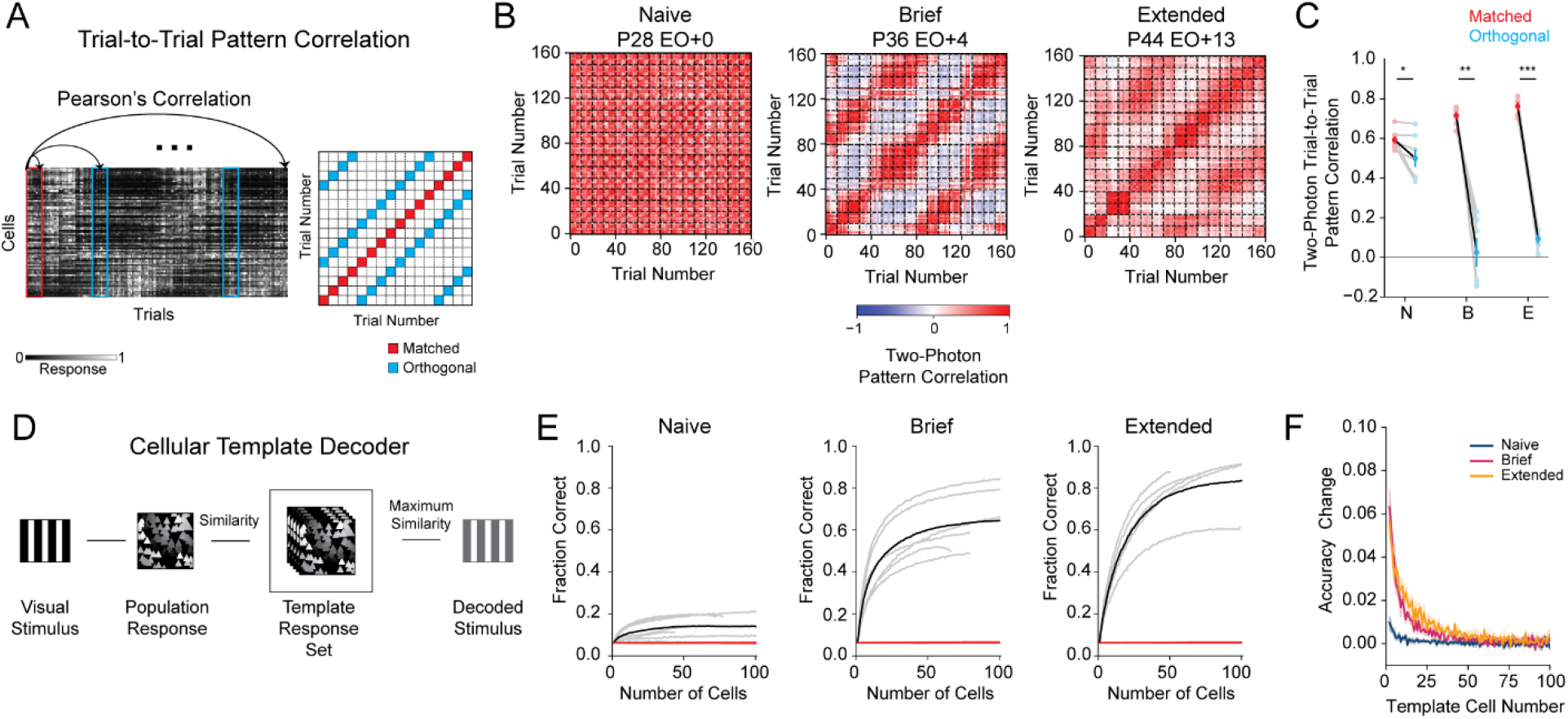
Population responses to orientation become more differentiable after visual experience. (A) Trial-to-trial pattern correlation matrices were generated by comparing population trial responses across stimulus conditions. (B) Example trial-to-trial pattern correlation matrices for a Naive (left), Brief (middle), and Extended (right) experience animal. (C) Comparisons of trial-to-trial pattern correlation for matched (red) and orthogonal (blue) stimuli across age groups (Naive: N (n=7), Brief: B (n=6), Extended: E (n=5)). (D) Schematic of the template decoding algorithm. (E) Decoding fraction versus number of cells used in the template decoder for Naive (left, n=7), Brief (middle, n=6), and Extended (right, n=5) animals. Results for individual animals (grey), grouped average (black), and shuffle (red). (F) Change in decoding accuracy versus the cell number in the decoder for Naive (blue, n=7), Brief (magenta, n-6), and Extended experience (gold, n=5). Shaded regions denote SEM. *: p<0.05, **: p<0.005, **: p<0.0005

To get a better understanding of the functional significance of these developmental changes in stimulus-evoked responses, we compared the accuracy of inhibitory networks from naive and experienced animals in discriminating orientations by using a template matching decoder^4,18^. Briefly, we generated a network pattern template set for each orientation from evoked activity patterns and compared them to individual trials to predict the stimulus orientation (Figure 2D). Across all cohorts, our template matching decoder predicted stimuli at a rate higher than chance (Figure 2E, Decoding Accuracy for 40 Cells compared to shuffle, Naive 0.136 ± 0.019, p=0.0094, n=7; Brief 0.578 ± 0.053, p=0.0002, n=6; Extended 0.720 ± 0.051, p=0.0002, n=5; chance level across all groups was 0.06, Mean ± SEM, Student’s t-test). Naive animals, however, for comparable numbers of cells decoded stimuli at a lower rate compared to Brief or Extended experience animals (decoding accuracy for 40 cells, Naive 0.136 ± 0.019 vs Brief 0.578 ± 0.053, p= 0.0006, n= 7 vs 6; Naive 0.136 ± 0.019 vs Extended 0.720 ± 0.051, p<0.0001, n= 7 vs 5; Mean ± SEM, bootstrap test). Consistent with these findings, when we assessed the change in decoding accuracy for templates including one cell compared to two, experienced animals showed a greater improvement in decoding accuracy (Figure 2F, Naive 0.010 ± 0.003 vs Brief 0.062 ± 0.008, p= 0.0002, n= 7 vs 6; Naive 0.010 ± 0.003 vs Extended 0.057 ± 0.005, p= 0.0004, n= 7 vs 5; Mean ± SEM, bootstrap test), suggesting individual cells carry more information about a stimulus later in development. In sum, these data demonstrate dramatic changes in the population response properties of GABA-INs after the onset of visual experience that endows them with a greater capacity to discriminate stimulus orientation.

### Clustered responses and modular patterns emerge after visual experience

We wondered how these changes in the responses of GABA-INs related to the modular organization of orientation preferences. Modular organization has been observed in both mature excitatory and inhibitory networks, as well as naive excitatory networks in the ferret^4,6,8^. To assess the state of the modular organization across development, we used widefield epifluorescence calcium imaging to visualize the response of GABA-INs at the millimeter scale (Figure 3A-B). In experienced animals, we observed clustered responses to oriented stimuli which gave rise to smoothly varying orientation preference maps (Figure 3B). Orientation preference maps, however, were not present in Naive animals and a smaller proportion of visually responsive pixels had significant orientation tuning (Figure 3D; Naive 0.32 ± 0.12 vs Brief 0.88 ± 0.02, p=0.0076, n= 5 vs 4; Naive 0.32 ± 0.12 vs Extended 0.94 ± 0.03, p= 0.0050, n= 5 vs 5; Mean ± SEM, bootstrap test).

**Figure 3.**
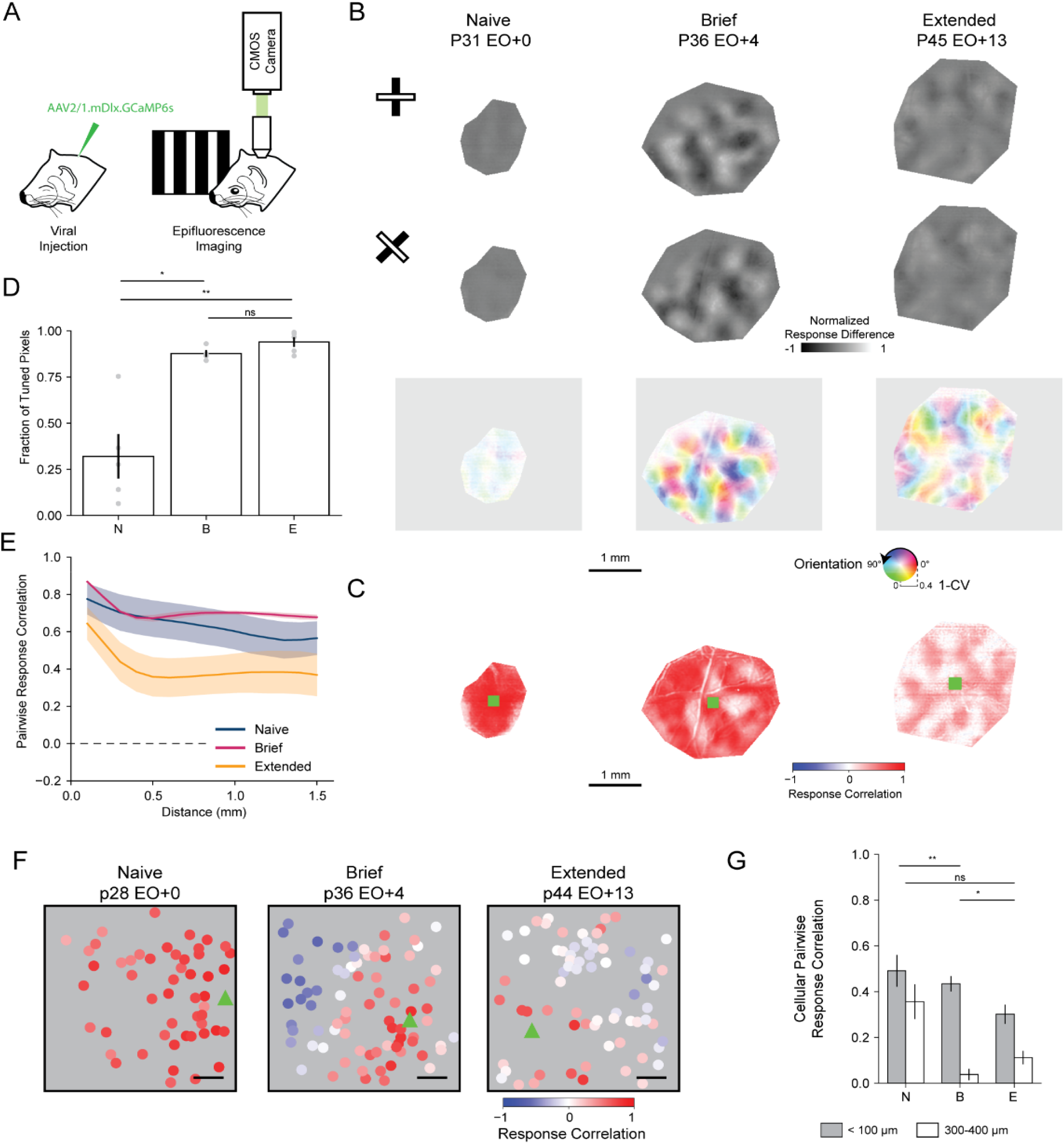
Clustered responses emerge across development in inhibitory networks. (A) Widefield epifluorescence calcium imaging of GCaMP6s under the mDlx enhancer in ferret visual cortex. (B) Difference Maps for orthogonal stimuli (top, middle) and orientation preference maps (bottom) for a Naive (left), Brief (Middle), and Extended (right) experience animal. (C) Example pairwise pixel response correlation maps for a Naive (left), Brief (Middle), and Extended (right) experience animal. Green squares identify the seed positions. (D) Fraction of pixels that are significantly orientation tuned compared to shuffle. Mean ± SEM. (E) Mean distance dependence of pairwise pixel response correlations for Naive (blue, n=5), Brief (magenta, n=4), and Extended (gold, n=5) experience animals. Shaded regions denote the SEM. (F) Example cellular pairwise correlation patterns for a Naive (left), Brief (middle), and Extended experience (right) animals. The seed cell is denoted by the green triangle and scale bar denotes 100 μm. (G) Comparisons of the average short-range (grey, < 100 μm) and long-range (white, 300-400 μm) cellular pairwise correlations for Naive (N, n=7), Brief (B, n=6), and Extended (E, n=5) experience animals. Mean ± SEM. *: p<0.05, **: p<0.005, ns: not significant

The absence of orientation preference maps in Naive animals, however, does not rule out the presence of modular patterns of activity in inhibitory networks. Indeed, Naive animals’ inhibitory network responses were poorly associated with oriented stimuli (see Figure 2B). Therefore, we employed a method for assessing modular network organization independent of stimulus identity. We computed correlation maps using pairwise pixel response correlations for drifting gratings to assess the structure of evoked responses independent of the visual stimuli presented. Consistent with our previous results, modular patterns were apparent in the response correlation maps for experienced animals (Figure 3C). Average correlation and distance displayed a complex relationship, deviating from linearity (Figure 3E, Brief linear regression R^2^= 0.33 ± 0.03, Extended linear regression R^2^= 0.35 ± 0.05). In contrast, we did not observe modular clustering in response correlation maps of Naive animals (Figure 3C). Furthermore, correlation strengths between pixels decreased almost linearly with distance (Figure 3E, Naive linear regression R^2^= 0.88 ± 0.06). Thus, naive inhibitory networks do not have any apparent modular organization at the millimeter-scale, and after a week of visual experience modular organization of responses and orientation preferences emerge.

While widefield imaging has shown that inhibitory networks do not display large-scale organization in the evoked responses in Naive animals, cellular-scale organization could still be present. For example, the lack of modular structure in Naive animals could arise from either the uniform activation of all cells or variable ‘salt and pepper’ activation of subsets of cells. To assess the cellular patterns of responses at a finer scale, we used two-photon imaging to measure the cellular pairwise correlations of evoked responses, independent of stimulus identity for short-range (< 100 μm) and long-range (300-400 μm) cell pairs. Across all age groups, cellular pairwise response correlations for short-range pairs were significantly higher than for long-range (Figure 3G, Naive short 0.49 ± 0.07 vs long 0.36 ± 0.08, p=0.0003, n= 7; Brief short 0.43 ± 0.03 vs long 0.04 ± 0.03, p =0.0009, n=6; Extended short 0.30 ± 0.04 vs long 0.11 ± 0.03, p=0.0004, n=5; Mean ± SEM, Student’s paired t-test), suggesting that salt and pepper response patterns are not typical. When comparing the relative difference between short- and long-range correlations across age, however, Naive animals had a significantly lower difference in clustering compared to animals with one week of visual experience (Figure 3G, Naive 0.14 ± 0.02 vs Brief 0.40 ± 0.06, p=0.0024, n= 7 vs 6, Mean ± SEM, bootstrap test). Importantly, cortical expansion cannot account for the increase in clustering between Naive and Brief experience animals, as it would predict a reduction in clustering. We also observed a significant reduction in the clustering between animals with Brief and Extended experience (Figure 3G, Brief 0.40 ± 0.06 vs Extended 0.19 ± 0.02, p=0.0106, bootstrap test), however, this could result from diversification of neuronal preferences along other relevant dimensions (such as spatial or temporal frequency) and/or cortical expansion. Our findings suggest that correlated responses in naive inhibitory networks are only modestly greater for neighboring cells than those at greater distances, and widespread, correlated activity across the network dominates responses. After eye opening, the correlation of neighboring cells remains high and long-range correlations decrease, resulting in patchy, clustered patterns of activity (Supplementary Figure 2).

### Development of orientation selectivity in inhibitory neurons depends on visual experience, but not restricted to the week after eye opening

Visual experience at eye opening is a necessary contributor to the normal maturation of excitatory networks and delayed experience onset results in deficits to orientation selectivity1,4. Therefore, we wondered whether inhibitory circuit development requires visual experience over the first week following normal eye opening. We performed binocular deprivation in a cohort of animals (BD) by eyelid suturing for 5-8 days, depriving them of structured visual experience. We then assessed the responses of cells to oriented stimuli after manually opening the eyes. Interneurons in BD animals had robust visually evoked activity (Figure 4A-C). The amplitude of these responses was not strongly dependent on the orientation or direction of the visual stimulus (Figure 4D, H). When compared to Brief experience animals, BD animals displayed significantly lower orientation selectivity (OSI BD 0.22 ± 0.04 vs Brief 0.54 ± 0.06, p= 0.009, n= 4 vs 6; Mean ± SEM, bootstrap test) and comparable direction selectivity (DSI BD 0.13 ± 0.03 vs Brief 0.19 ± 0.03, p=0.5196, n= 4 vs 6; Mean ± SEM, bootstrap test). In addition, BD animals showed a higher degree of variability in responses (VI BD 0.75 ± 0.01 vs Brief 0.48 ± 0.04, p= 0.0038, n= 4 vs 6; Mean ± SEM, bootstrap test). Together these data suggest that the functional properties of inhibitory neurons require visual experience to reach mature levels.

**Figure 4.**
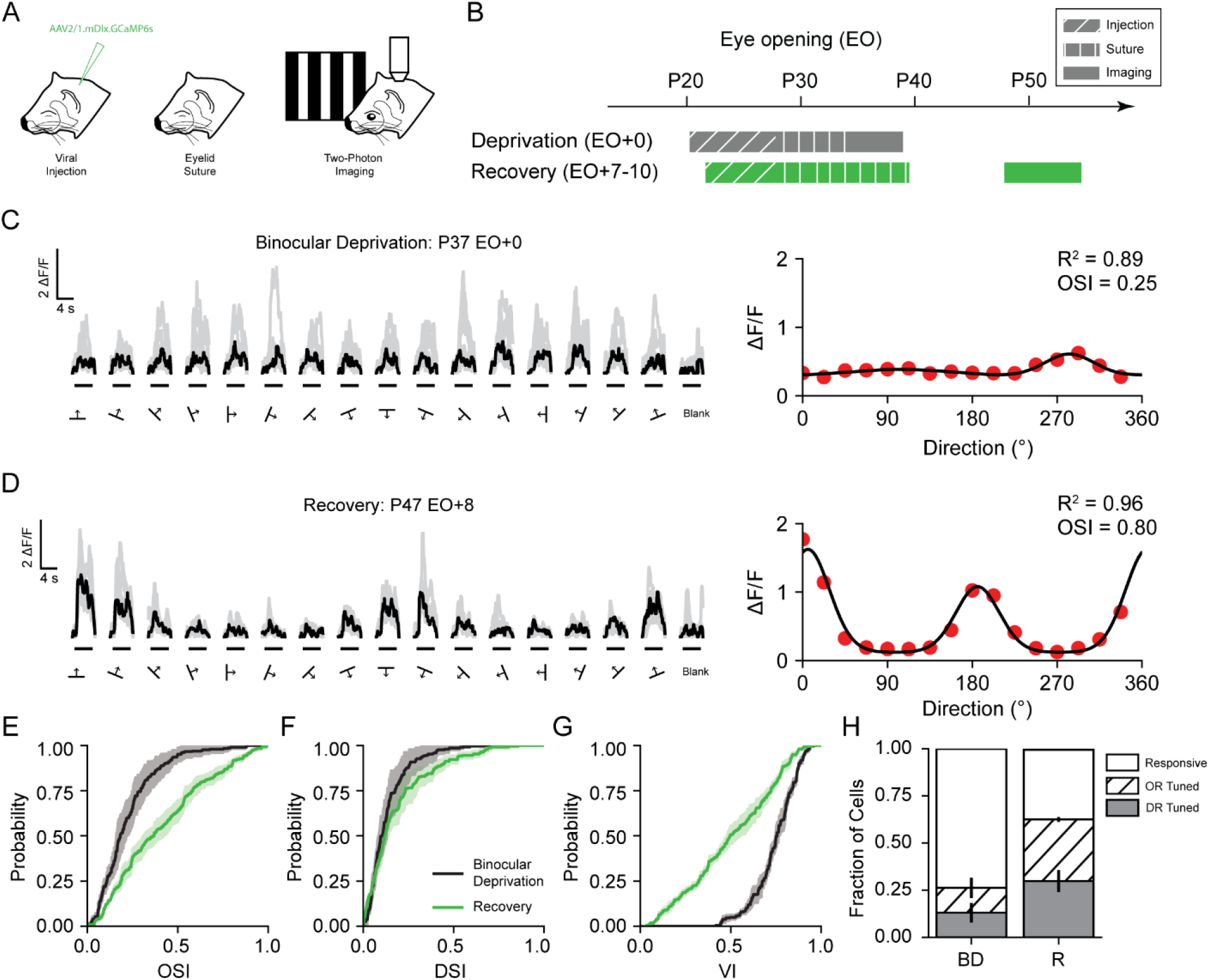
Development of orientation preference in interneurons requires visual experience. (A) Two-photon imaging of viral expression of GCaMP6s under the mDlx enhancer in ferret visual cortex after binocular eyelid suture (B) Two cohorts, Binocular Deprivation (EO+0, 5-8 days of deprivation) and Recovery (EO+8-15, 9 days of deprivation) were evaluated for GABA-IN responses to oriented stimuli. (C) Example cellular response to drifting gratings (left, black traces denote median response and gray traces denote single trials) and response tuning curve (right) for a binocularly deprived animal. (D) Example cellular response to drifting gratings (left) and response tuning curve (right) for a recovery animal. (E-G) Cumulative distributions of orientation selectivity index (E), Cumulative distributions of direction selectivity (F), and cumulative distributions of variability index (G) for BD (black, n=4) and Recovery (green, n=4) animals. Shaded regions denote SEM. (H) Fraction of cells that are responsive (white), orientation selective (hatched), and direction selective (gray) for Binocular Deprivation (BD, n=4) and Recovery (R, n=4) animals. Error bars denote SEM.

To test the necessity of visual experience over the first week following normal eye opening, we performed binocular eyelid suturing for 9 days, manually opened the eyes, and then assessed visual responses after 8-15 days of visual experience (Figure 4A-B, Recovery). Interneurons in recovery animals responded robustly to oriented stimuli (Figure 4D). BD animals had significantly lower orientation selectivity than Recovery animals (Figure 4E, OSI BD 0.22 ± 0.04 vs Recovery 0.40 ± 0.03, p= 0.0170, n= 4 vs 4, bootstrap test). Direction selectivity of orientation selective cells, however, did not significantly change (Figure 4F, DSI BD 0.13 ± 0.03 vs Recovery 0.18 ± 0.03, p= 0.2350, n = 4 vs 4, bootstrap test). In addition, cellular response variability decreased with experience (Figure 4G, VI BD 0.75 ± 0.01 vs Recovery 0.50 ± 0.02, p= 0.0042, n= 4 vs 4, bootstrap test). When comparing Recovery and Extended experience animals, however, we found no significant difference in orientation selectivity, direction selectivity, or cellular variability (OSI, Recovery 0.41 ± 0.03 vs Extended 0.52 ± 0.05, p= 0.0650; DSI, Recovery 0.18 ± 0.03 vs Extended 0.24 ± 0.03, p = 0.1216; VI, Recovery 0.50 ± 0.02 vs Extended 0.48 ± 0.02, p= 0.3648; n = 4 vs 5, bootstrap test). In sum, these data suggest that the maturation of orientation selectivity and reduction in cellular variability of GABA-INs depend on visual experience but can occur beyond the first week following normal eye opening.

### Development of orientation-specific population responses requires visual experience

We next assessed the correspondence of evoked population patterns to presented stimuli, by assessing the trial-to-trial pattern correlation for matched and orthogonal orientations (Figure 5A-B). BD animals displayed a high degree of similarity in response pattern correlations for matched and orthogonal orientations (Figure 5B, BD Matched 0.59 ± 0.02 vs Orthogonal 0.49 ± 0.05, p= 0.0718, n=4, Mean ± SEM, Student’s paired t-test). The response correlations for matched and orthogonal orientations were not significantly different from Naive animals (BD matched 0.59 ± 0.03 vs Naive 0.59 ± 0.03, p= 0.9562; BD orthogonal 0.49 ± 0.05 vs Naive orthogonal 0.50 ± 0.04, p= 0.8114, n=4 vs 7, Mean ± SEM, Student’s paired t-test). Thus, inhibitory networks in BD animals lack a population-level representation of orientation, and visual experience is necessary for the formation of orientation-specific responses.

**Figure 5.**
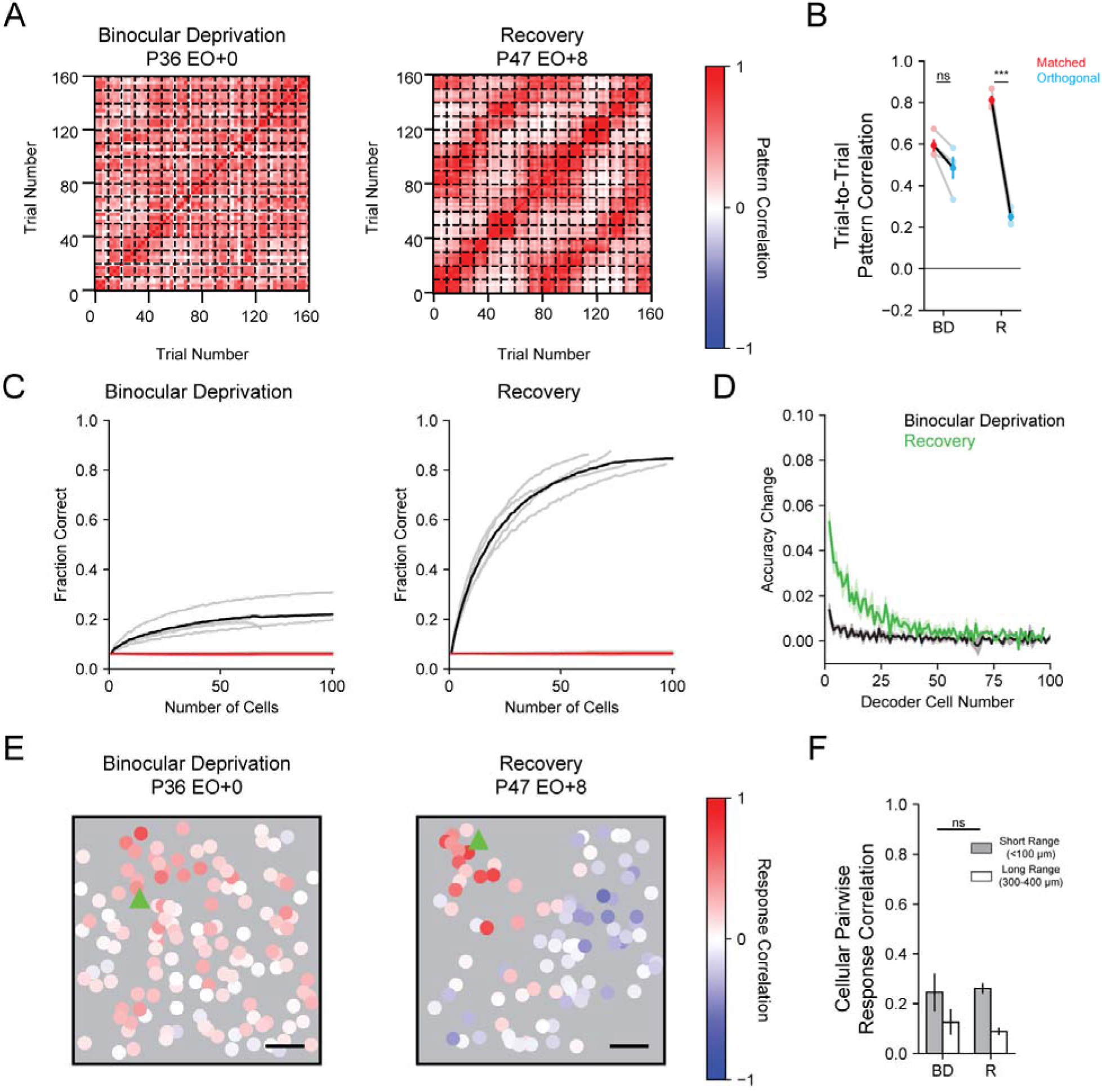
Reduction in spatially uniform responses occurs without visual experience. (A) Trial-to-trial pattern correlations for a Binocular Deprivation (left) and Recovery (right) animal. Trials are grouped by stimulus direction and dotted lines denote groups. (B) Comparison of matched (red) and orthogonal (blue) trial-to-trial pattern correlations for Binocular Deprivation (n=4) and Recovery (n=4) animals. Individual animals are shown as gray and error bars denote SEM. (C) Template decoding accuracy for Binocular Deprivation (n=4) and Recovery (n=4) animals for different template sizes. Red line indicates chance decoding levels, black shows average decoding level, and gray lines denote individual animals. (D) Change in decoder accuracy for the nth cell included in the template decoder for Binocular Deprivation (black, n=4) and Recovery (green, n=4) animals. Shaded regions denote SEM. (E) Example cellular pairwise correlations for Binocular Deprivation (left) and Recovery (right) animals. The seed cell is denoted by the green triangle and scale bar denotes 100 μm. (F) Binocular Deprivation (n=4) and Recovery (n=4) differences of cellular pairwise response correlations for short-(< 100 μm) and long-range (300-400 μm) pair are similar. ***: p<0.0005, ns: not significant

If the development of orientation-specific responses requires visual experience, is delayed visual experience sufficient to recover orientation-specific responses? In Recovery animals, response pattern correlations were higher for matched stimuli (Matched Stimuli Pattern Correlation, Recovery 0.81 ± 0.02 vs BD 0.59 ± 0.03, p= 0.0064, n= 4 vs 4; Mean ± SEM, bootstrap test) and lower for orthogonal stimuli (Orthogonal Stimuli Pattern Correlation, Recovery 0.25 ± 0.02 vs BD 0.49 ± 0.05, p= 0.0108, n= 4 vs 4; Mean ± SEM, bootstrap test). We observed similar differences when comparing Recovery and Naive animals (Recovery matched 0.81 ± 0.02 vs Naive 0.59 ± 0.02, p= 0.0014; Recovery orthogonal 0.2503 ± 0.02 vs Naive 0.50 ± 0.04, p= 0.0048; n= 7 vs 4, Mean ± SEM, bootstrap test). Age matched animals with normal visual experience also displayed higher similarity of response patterns for matched stimuli, and lower similarity for orthogonal stimuli (matched Recovery 0.81 ± 0.02 vs Extended 0.76 ± 0.02, p= 0.1118; orthogonal Recovery 0.25 ± 0.02 vs Extended 0.09 ± 0.02, p=0.0076; n= 5 vs 4, Mean ± SEM, bootstrap test). These data suggest that orientation-related inhibitory network pattern development is resilient to the delayed onset of visual experience.

We further assessed the relationship of stimulus-evoked responses to orientation by testing a template decoding algorithm for BD and Recovery animals (Figure 5C). Consistent with our trial-to-trial pattern correlation findings, BD animals significantly underperformed as compared to Recovery animals (Decoding Accuracy for 40 cell template BD 0.185 ± 0.023 vs Recovery 0.713 ± 0.031, p=0.8950, n= 4 vs 4, Mean ± SEM, bootstrap test). Recovery animals also showed marked improvement in decoding of orientation even for templates using small numbers of cells (Change in decoding accuracy for 2^nd^ cell, BD 0.014 ± 0.03 vs Recovery 0.053 ± 0.005, p= 0.0038, n= 4 vs 4, Mean ± SEM, bootstrap test). Thus, visual experience is necessary for the development of orientation-related responses of GABA-INs, improving the discriminability of orthogonal stimuli responses and improving response reliability.

While Recovery animals showed improvement in orientation-related response discriminability, it remains possible that the inhibitory networks of these animals may have an impaired encoding of orientations. Recovery animals, however, did not significantly underperform when compared to age-matched animals with normal experience (Decoding Accuracy for 40 cell template Recovery 0.713 ± 0.031 vs Extended 0.720 ± 0.051, p= 0.8950, n= 5 vs 4; Mean ± SEM, bootstrap), and the change in decoding accuracy for small numbers of cells was also comparable (Figure 5D, Change in decoding accuracy for 2^nd^ cell, Recovery 0.053 ± 0.005 vs Extended 0.057 ± 0.005, p=0.5606, n= 5 vs 4, Mean ± SEM, bootstrap test) suggesting that there is not a strict critical period for the experience-dependent development of orientation-specific population responses for inhibitory networks.

### Developmental reduction in non-modular responses does not require visual experience

These results indicate that experience plays a key role in the development of orientation selective responses in GABA-INs, and one might expect that experience plays a similar role in the development of modular spatial organization for GABA-INs. First, we assessed the spatial organization of response correlations (Figure 5E), and found that both BD and Recovery animals demonstrated significant higher correlations for short-range cell pairs (<100 μm) compared to long-range cell pairs (300-400 μm) (Figure 5F, BD short 0.25 ± 0.08 vs long 0.13 ± 0.05, p= 0.0357, n= 4; Recovery short 0.26 ± 0.02 vs long 0.09 ± 0.02, p= 0.0028, n= 4; Mean ± SEM, Student’s paired t-test), and the difference in correlations for short- and long-range cell pairs were similar when compared to animals with normal extended experience (BD 0.12 ± 0.03 vs Extended 0.19 ± 0.02, p=0.0630, n= 4 vs 4, Mean ± SEM, bootstrap test). Importantly, correlations for both short- and long-range cell pairs were systematically lower for BD animals when compared to Naive animals (Naive short 0.49 ± 0.07 vs BD short 0.25 ± 0.08, p=0.0392; Naive long 0.36 ±0.08 vs BD long 0.13 ± 0.05, p= 0.0448; n= 4 vs 7, Mean ± SEM, bootstrap test), suggesting that the spatially uniform responses that dominate responses in Naive animals were greatly reduced in BD animals. Furthermore, in Recovery animals, subsequent experience did not affect the difference in short- and long-range correlations (BD 0.12 ± 0.03 vs Recovery 0.17 ± 0.02, p= 0.1572; n= 4 vs 4, Mean ± SEM, bootstrap test), suggesting delayed visual experience does not alter the degree to which responses to oriented stimuli cluster.

How could clustered responses in BD animals coexist with poor orientation tuning? One possibility is pattern variability-wherein visual stimuli inconsistently recruit modular patterns of activity. The result of pattern variability is that on a trial-to-trial basis modular patterns are observable, but stimulus-averaged responses would lack modular structure. Consistent with such a mechanism, we observe lower pattern correlations for matched orientations (Figure 5B) and a diversity of patterns for trial activity (Supplementary Figure 3) in BD animals compared to Recovery animals. Together, our findings demonstrate that the reduction of spatially uniform responses and subsequent unmasking of clustered activity in inhibitory networks can develop in an experience-independent manner over the developmental period spanning the first week of normal visual experience, while the development of consistent, orientation associated modular responses requires visual experience.

## Discussion

In summary, we have demonstrated that before visual experience, interneurons in the ferret visual cortex show poor cellular orientation tuning due to highly variable cellular responses. These variable responses manifest as a global pattern of activity that recruits all interneurons independent of the orientation of visual stimuli. As little as a week of visual experience, however, suffices to drive the development of strong orientation tuned responses, reduction in uniform activity, and the consistent association of modular patterns to stimuli. Furthermore, interneurons are resilient to delayed experience onset- maintaining the ability to develop strong orientation tuning and orientation associated modular network responses beyond the first week of normal visual experience. Finally, the development of modular inhibitory responses does not require visual experience as we observed modular response patterns in animals after a week of binocular deprivation. However, the association of these patterns to oriented stimuli requires visual experience, as binocularly deprived animals displayed deficits in orientation selectivity and network pattern differentiability.

In the mouse visual cortex, it has been hypothesized that the broadening of tuned inhibition is a critical factor gating the plasticity of the monocular matching of orientation preferences of excitatory cells^15^. However, the murine visual cortex has salt and pepper organization of orientation preferences and low orientation selectivity in mature interneurons^13,14,19^. Our observations in the modularly organized ferret visual cortex demonstrate that over a comparable developmental period, when excitatory neurons match monocular and binocular orientation preference^4^, GABA-INs are gaining orientation selectivity. Thus, our findings challenge the generality of experience-dependent broadening of tuned inhibition playing a role in initiating the critical period for excitatory binocular plasticity in excitatory networks of the visual cortex. Studies on inhibitory development to date have predominantly focused on the impact of changes in the strength of inhibition, independent of functional properties, as an important factor gating critical period plasticity. Modulating the strength of GABAergic inhibition can alter the timing of the critical period^12,20^ and development of modular cortex^21^, and transplantation of GABA-INs can rescue critical period-like plasticity in adults^22–24^. In addition, early dysfunction of GABA-INs can lead to impairment of visual circuits^10,11,25^. In contrast, our findings highlight that beyond changes in the strength of inhibition, functional properties of GABA-INs also shift after eye opening, and these changes in functional properties could contribute to the normal development of the excitatory network.

The influence of GABA-INs on orientation tuning in adult animals has been an area of great interest. Early studies suggested that lateral-inhibition shaped orientation tuning by suppressing responses to orthogonal orientations^26,27^. Recent investigations, however, have consistently found weak inhibition for orthogonal orientations- instead finding either untuned or co-tuned inhibition^28–30^, suggesting that lateral-inhibition is not a requirement for orientation preferences of excitatory neurons in mature animals. While lateral-inhibition is not a key contributor to mature orientation preferences, there remained the possibility that a transient phase of lateral-inhibition early in development could play a role in establishing orientation preferences. Our observations that before the onset of sensory experience, GABA-INs in the modular visual cortex of ferret are not well orientation tuned are inconsistent with this possibility. Therefore, excitatory orientation preferences are likely to result from processes that do not require tuned inhibition.

Intrinsic unreliability of sensory input to the visual cortex (sensory noise) could account for the variability in response patterns. One strategy for handling sensory noise is distributing information about sensory stimuli across populations of neurons^32^, and in visual cortex, this distributed representation results in orientation preference maps^2,6^. In both Naive and BD animals, the accuracy of predicting stimulus identity based on population response is impaired because of an underdeveloped representation of orientation. Unlike Naive animals, however, the resulting patterns of activity in BD animals display clustered responses that are poorly associated with the presented visual stimuli. We have also observed similar pattern variability in excitatory networks of visual cortex before eye opening, where clustered responses are apparent^33^. Together these findings suggest that the underlying synaptic connectivity of cortical neurons can limit the co-activation of populations of cells, and modular activity on a trial-to-trial basis can occur independent of orientation tuning if feedforward sensory inputs unreliably recruit patterns of activity.

GABA-INs encompass a diversity of cells across different lamina, with different genetic markers, cellular morphology, and synaptic targets in the visual cortex. For example, interneurons expressing parvalbumin make perisomatic and axonal contacts, while interneurons expressing the peptide somatostatin have synaptic contacts biased toward the distal dendrites suggesting subtypes of interneurons may play different roles in shaping cortical activity^34–36^. Additionally, optogenetic modulation of interneuron activity has subtype-specific effects on visually evoked activity^37–41^, and subtype-specific synaptic changes coinciding with the onset of visual experience have also been observed^42^. Viral expression using the mDlx enhancer labels a broad diversity of interneurons including cells that express somatostatin and parvalbumin, and differences in functional properties for these inhibitory cell types have been observed in mature ferrets^8,17^. Because of limitations in identifying cells based on molecular markers early in development^43^, we pooled all GABA-INs. Importantly, we did not observe significant heterogeneity in cellular responses of Naive animals, suggesting that most interneurons do not exhibit modular responses or orientation tuning early in development. Indeed, novel methods targeting interneuron subtypes open up the possibility of assessing subtype-specific developmental trajectories of GABA-INs^44^.

What contributes to the development of orientation preferences in GABA-INs? We observe that evoked responses show nearly uniform participation of interneurons in naive animals in primary visual cortex. Thus, it remains likely that a strong shared sensory drive recruits the entire inhibitory network. One possibility is that uniform responses result from lateral inputs from neighboring layer II/III neurons that have a diverse spectrum of orientation preferences. In aggregate, these inputs could drive the non-orientation-specific responses seen early in development, and the subsequent refinement of the orientation preferences of these inputs could drive the gain of orientation tuning in interneurons. Alternatively, untuned responses in interneurons could arise from strong untuned inputs, possibly originating from the LGN or layer IV that occlude well-organized lateral inputs from layer II/III during visual stimulation. The subsequent loss of these untuned inputs could unmask orientation tuned responses in interneurons after eye opening. Such a mechanism would be consistent with recent observations of spontaneous modular inhibitory activity before the onset of vision^45^, and anatomical studies that have established reduction in thalamocortical inputs to cortical interneurons across early development and the classical ocular dominance critical period^46–48^. Further investigation into the synaptic inputs to GABA-INs across development is necessary to clarify these possibilities. In conclusion, our findings, highlight that interneurons undergo a parallel, delayed process in which experience-independent mechanisms reduce widespread, network responses, and experience-dependent processes drive the development of orientation-specific responses, resulting in modular, functional networks of GABA-INs.

## Methods

### Animals

All experimental procedures were approved by the Max Planck Florida Institute for Neuroscience Institutional Animal Care and Use committee and were performed in accordance with guidelines from the U.S. National Institute of health. Juvenile female ferrets (*Mustela putorius furo*, Marshal Farms) co-housed with jills on a 16-h light/8-h dark cycle. No *a priori* sample size estimation was performed, but sample sizes are comparable to other studies which performed *in vivo* imaging.

### Viral Injections and Eyelid Suturing

We expressed GCaMP6s by microinjection of a custom-made virus AAV2/1-mDlx-GCaMP6s^28^ (Vigene), at 6-21 days before imaging experiments. AAV2/1-mDlx-GCaMP6s was derived from pAAV-mDlx-GCaMP6f-Fishell-2, which was a gift from Gordon Fishell (Addgene plasmid # 83899; http://n2t.net/addgene:83899; RRID:Addgene_83899), and has previously been shown to specifically target a diversity of interneurons subtypes in the ferret visual cortex^8,17^. Anesthesia induction was performed using either ketamine (50mg/kg, IM) and/or isoflurane (1-3%) delivered in O2 and then maintained with isoflurane (1-2%). Atropine (0.2mg/kg, IM) was administered to reduce secretions, while Buprenorphine (0.01mg/kg, IM) and a 1:1 mixture of lidocaine and bupivacaine (injected directly into the scalp) were administered as analgesics. Animal temperatures were maintained at 37°C using a homoeothermic heating blanket.

Animals were also mechanically ventilated and both heart rate and end-tidal CO_2_ were monitored throughout the surgery. Under aseptic surgical technique, a small craniotomy was made over visual cortex 6.5-7 mm lateral and 2mm anterior to lambda. Approximately 1μL of virus was pressure infused into the cortex through a pulled glass pipette across two depths (~200μm and 400μm below the surface). For eyelid suturing, eyelids were binocularly sutured during a short aseptic technique between P26-30. Eyelid sutures were monitored daily until they were removed for imaging experiments as noted in the main text.

### Cranial Window and Acute Imaging

All animals were anesthetized and prepared for surgery as described above. In addition, an IV catheter was placed in the cephalic vein. Preparation for cranial windows and acute imaging was performed as previously described^4^. Briefly, a custom-designed metal headplate (8 mm DIA) was implanted over the injected region and secured with MetaBond (Parkell Inc.). Then a craniotomy and durotomy were performed, exposing the underlying brain. The brain was then stabilized with a custom-designed titanium metal cannula (4.5 mm DIA, 1.5 mm height), adhered to a 4mm coverslip (Warner Instruments) with optical glue (#71, Norland Products Inc.). Finally, the headplate was hermetically sealed with a stainless-steel retaining ring (5/16” internal retaining ring, McMaster-Carr) and glue (Krazy Glue).

For all imaging experiments, either eyelid sutures were removed, or the eyelids were separated where applicable to ensure visual stimulation was presented to open eyes. Phenylephrin (1.25-5%) and tropicamide (0.5%) were applied to the eyes to retract the nictating membrane and dilate the pupils. Prior to imaging, isoflurane levels were reduced from a surgical plan to ~1-1.5%. After a stable, anesthetic baseline was established for 30 minutes, animals were paralyzed with pancaronium bromide (0.1-1 mg/kg/hr).

### Calcium Imaging Experiments

All imaging experiments were preformed using a B-Scope microscope (Thorlabs) and began immediately after cranial window surgery. Two-photon experiments of GCaMP6s used a Mai-Tai DeepSee laser (Spectra Physics) at 910 nm for excitation. The B-Scope microscope was controlled by ScanImage (Vidreo Technologies) in a resonant-galvo configuration with images acquired at 512×512 pixels. Multi-plane images were sequentially collected from one or four imaging planes using a piezoelectric actuator for an effective frame rate of 30 or 6 Hz respectively. Images were acquired at 2x zoom through a 16x water-immersion objective (Nikon, LWD 16X W/0.8 NA) yielding a field-of-view of ~0.7mm square (1.36 μm/pixel). All two-photon imaging planes were restricted to 100-250 μm from the cortical surface in order to assess responses in layer 2/3. For widefield epifluorescence imaging experiments, Zyla 5.5 sCMOS camera (Andor) controlled by μManager^49^. Images were acquired at 15 Hz using a 4×4 binning to yield 640×540 pixel images through a 4x air objective (Olympus, UPlanFL N 4x/0.13 NA).

### Visual Stimulation

Visual stimuli were presented on an LCD screen played approximately 25 cm from the eyes using PsychoPy^50^. To evoke orientation-specific responses, full-field square gratings at 100% contrast drifting at 1 Hz were presented in 16 directions. In addition, “blank” stimuli of 0% contrast were also presented to establish threshold for visual responsiveness. All stimuli were randomly interleaved and presented for 4 seconds follow by 6 seconds of gray screen.

### Analysis

All data analysis was performed using custom written scripts in either Matlab, ImageJ, or Python (Python 3.6). For both two-photon and widefield epifluorescence imaging data we corrected brain movement during imaging by maximizing phase correlation to a common reference frame. Statistical testing was performed by either bootstrap resampling or Student’s paired t-test, on a per animal basis unless otherwise noted in the text. For bootstrap resampling, 10000 resamplings with replacement were made. Two-tailed assessment of significance were made by comparing observed group averages to the 95^th^ percentile of the extremes (highest 97.5^th^ percentile or lowest 2.5th percentile) of the surrogate distributions. We chose to use bootstrap statistical testing for unpaired data analysis, as it is not dependent on assumptions about the underlying distributions present in our data.

#### Two-Photon Analysis

To identify cellular ROIs in two-photon imaging, custom software in ImageJ (Cell Magic Wand^8^) was used. ΔF/F for each ROI was computed by calculating the F_0_ using a rolling window (60 seconds) rank-order filter to the raw fluorescence (20^th^ percentile). Stimulus-evoked responses were computed by subtracting the mean pre-stimulus ΔF/F (1 second before stimulus onset) from the mean stimulus-evoked response over the entire stimulus period (4 s). Visually responsive cells were defined as cells where the maximum stimulus-evoked response was at least two standard deviations larger than the blank stimulus responses.

Since a fraction of interneurons are expected to have direction selective responses, all analyses were performed in direction space in order to minimize any impact of direction selectivity on both cellular and population responses. To assess orientation preference two von Mises functions, constrained to be 180° out of phase, were fit to the trial-median responses to stimulus directions. Orientation preference was assessed as the orientation for the peak response. Cells were considered well-tuned if the R^2^ of the fit were >0.6. Orientation selectivity was computed using the fit orientation tuning curves:

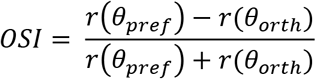

Where *r(θ)* is the fit tuning curve response for either the preferred direction or the orthogonal directions. Similarly, direction selectivity was computed:

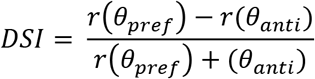

To test for significant orientation and direction tuning, we computed orientation tuning curve fits, OSI, and DSI for 1000 bootstrap with replacement samplings of trial evoked responses. Cell were deemed significantly orientation or direction tuned if the observed OSI or DSI was larger than the 95^th^ percentile of computed bootstrap OSI or DSI respectively. To assess the variability of cells, trial responses within stimuli were resampled 1000 times to compute resampled tuning curves (*T_n_*). The VI was then defined as:

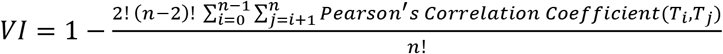

Cellular pairwise correlation maps were generated by computing the Pearson’s correlation of cellular responses to square wave drifting grating, independent of the stimulus identity. Short- and long-range cellular correlations were computed by taking the average pairwise correlation for cells within 100 μm and 300-400 μm respectively. Trial-to-trial pattern correlations were computed by taking all cells within a field-of-view and computing Pearson’s correlation coefficients across trial pairs. Matched and orthogonal trial-to-trial correlations were computed as the mean correlation for matched and orthogonal stimuli. We chose to perform these comparisons in direction space to minimize differences in evoked patterns due to increasing direction selectivity across development.

Finally, we implemented a normalized template matching algorithm as previously described^4^. Briefly, template population responses for each stimulus condition were generated by computing the median response using half the trials across all direction conditions. We assessed the performance of the template decoder by shuffling trials and increasing the number of cells included in the templates to generate template decoding accuracy versus template size. To assess the overall performance of the template decoders, we chose to use 40 cell templates, as the accuracy across all conditions appeared to approach peak accuracy near template size. To assess chance decoding rates, we generated templates using shuffled trials and assessed the decoding accuracy of a shuffled test data set (n=1000).

#### Widefield Epifluorescence Analysis

ROI segmentation was performed in widefield epifluorescence imaging by manually drawing around cortical regions where robust visual evoked activity was observed. For analysis, all images were spatially downsampled by a factor of 2x to yield 320×270 pixels at a spatial resolution of 11.63 μm/pixel. Slow drifts in fluorescence intensity were eliminated by calculating the ΔF/F. For each pixel, the baseline F0 was calculated by applying a rank-order filter to the raw fluorescence trace (10th percentile) with a rolling time window of 60 seconds.

Preferred orientation was computed as the vector sum of the average response for each orientation:

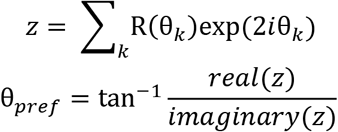

where R(θ) is the median response for a given orientation θ. To evaluate orientation selective pixels, we computed 1-CV metrics using the following formula:

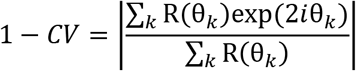

Significantly tuned pixels were assessed by computing 1-CV for 1000 bootstrapped with replacement re-samplings. Pixels were deemed significant if the observed 1-CV greater than 95% of the resampled 1-CVs. Correlation maps were computed using seed pixels and calculating the Pearson’s correlation coefficient for all other pixels with the ROI. Correlations were then binned by pixel distances to generate correlation as a function of distance plots, and linearity (R^2^) was assessed by performing ordinary least squares linear progression. Finally, trial-to-trial correlation matrices were generated by, by computing the Pearson’s correlation coefficient for all possible combinations of trials. Trial-to-trial correlations for matched and orthogonal stimuli were computed by taking the average correlation for sets of trials with matched stimuli and orthogonal stimuli in direction space.

#### Statistics

Our two photon analyses were based on 781 cells from 7 Naive animals, 1185 cells from 6 Brief experience animals, 541 cells from 5 Extended experience animals, 394 cells from 4 Binocular Deprivation animals, and 310 cells from 4 Recovery animals. Widefield calcium imaging was based on 5 Naive animals, 4 Brief experience animals, and 5 Extended experience animals. Unless otherwise noted, hypothesis testing was performed using a bootstrapping test on a per animal basis. We deliberately chose to use resampling methods for hypothesis testing, rather than relying on standard parametric or non-parametric tests, to avoid confounds that could arise from making assumptions on the underlying sampling distributions present in our data. Statistical significance was assessed by ranking if the observed difference between group averages was more extreme than the 95th percentile from a surrogate distribution, where the null hypothesis is true. All statistical tests were two-tail, except when assessing if the performance of the template matching decoder was above chance-levels (one-tail). For each hypothesis test, we produced a relevant surrogate distribution by pooling data between groups, bootstrapped new group averages from the pooled data using replacement (n=1,000), and then computed a distribution of differences between surrogate group averages. Paired tests for pattern correlation and response clustering were performed using a two-tailed paired Student’s t-test.

## Data Availability

Data are available from the corresponding author upon reasonable request.

## Code Availability

Code are available from the corresponding author upon reasonable request

## Author Contributions

All authors designed the study, analyzed the results, and wrote the paper. J.C. performed the widefield and two-photon calcium imaging.

## Acknowledgements

We would like to thank N. Shultz, R. Satterfield, J. Kerr, G. Kreal, and D. Whitney for technical assistance, as well as members of the Fitzpatrick laboratory for helpful discussions. This research was supported by US National Institutes of health grants EY011488 and EY026273 and the Max Planck Florida Institute for Neuroscience.

## Declaration of Interests

The authors declare no competing interests.

## Notes

### Competing Interest Statement

The authors have declared no competing interest.

