## Supplemental Figures for "Development of Visual Response Selectivity in Cortical GABAergic Interneurons"

### Supplementary Figures

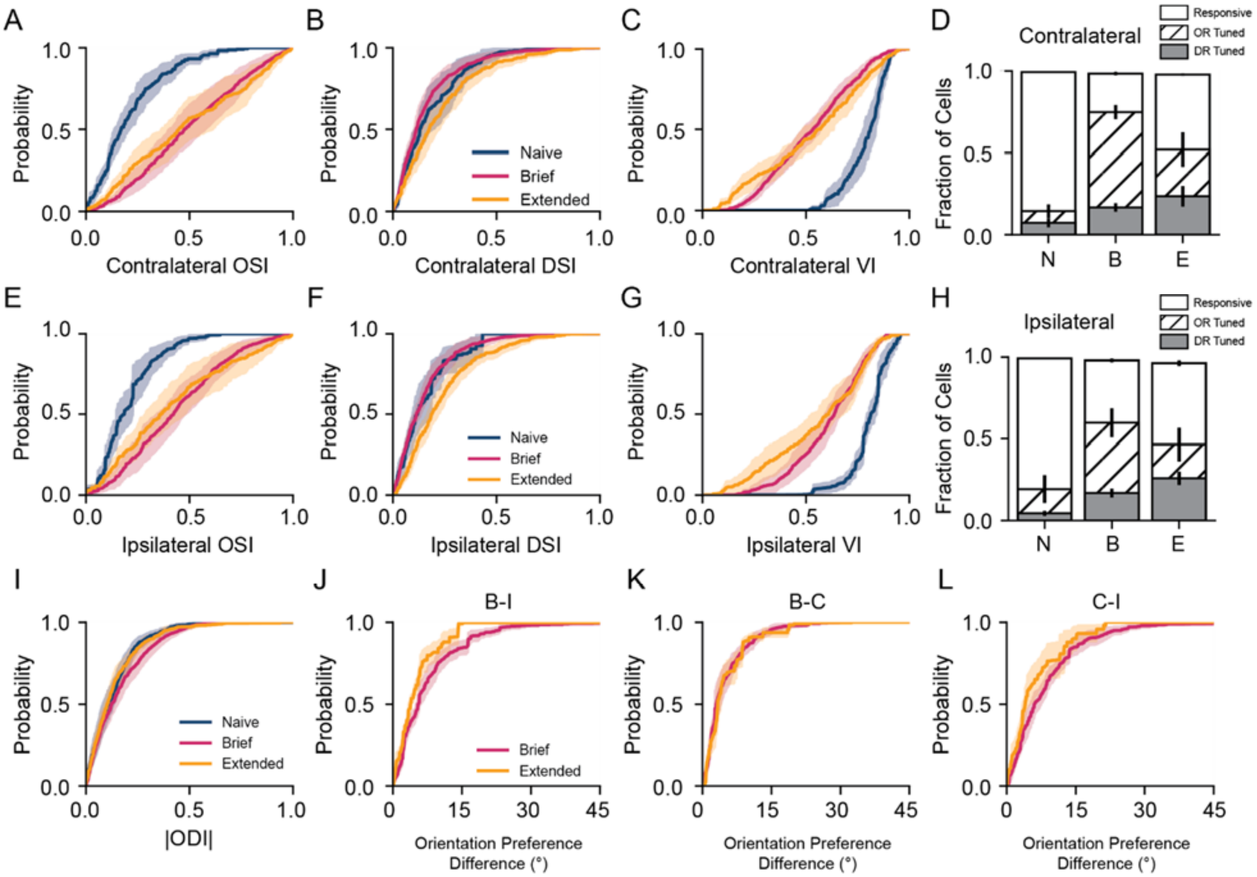


**Supplementary Figure 1 Binocular orientation binocular preferences develop in an aligned manner.** (A-B) Orientation selectivity index (A), direction selectivity index (B), and variability index (C) for contralateral stimulus presentation for Naive (blue), Brief (Magenta), and Extended (gold) experience. Mean ± SEM. (D) Fractions of cells that are responsive (open), significantly orientation tuned (hatched), or significantly direction tuned (gray) for Naive (N), Brief experience (B), and Extended (E) experience. Error bars denote SEM. (E-G) Same as A-B but for ipsilateral stimulus presentation. (H) same as D but for ipsilateral stimulus presentation. (I) Cumulative plot of monocularity (|ODI|) for Naive (blue), Brief (magenta), and Extended (gold) experience. (J) Cumulative plot of the difference in preferred orientation for binocular versus ipsilateral stimulus presentation. (K) Cumulative plot of the difference in preferred orientation for binocular versus contralateral stimulus presentation. (L) Cumulative plot of the difference in preferred orientation for contralateral versus ipsilateral stimulus presentation. Mean ± SEM. For all plots Naive (n=7), Brief (n=6), and Extended (n=7)

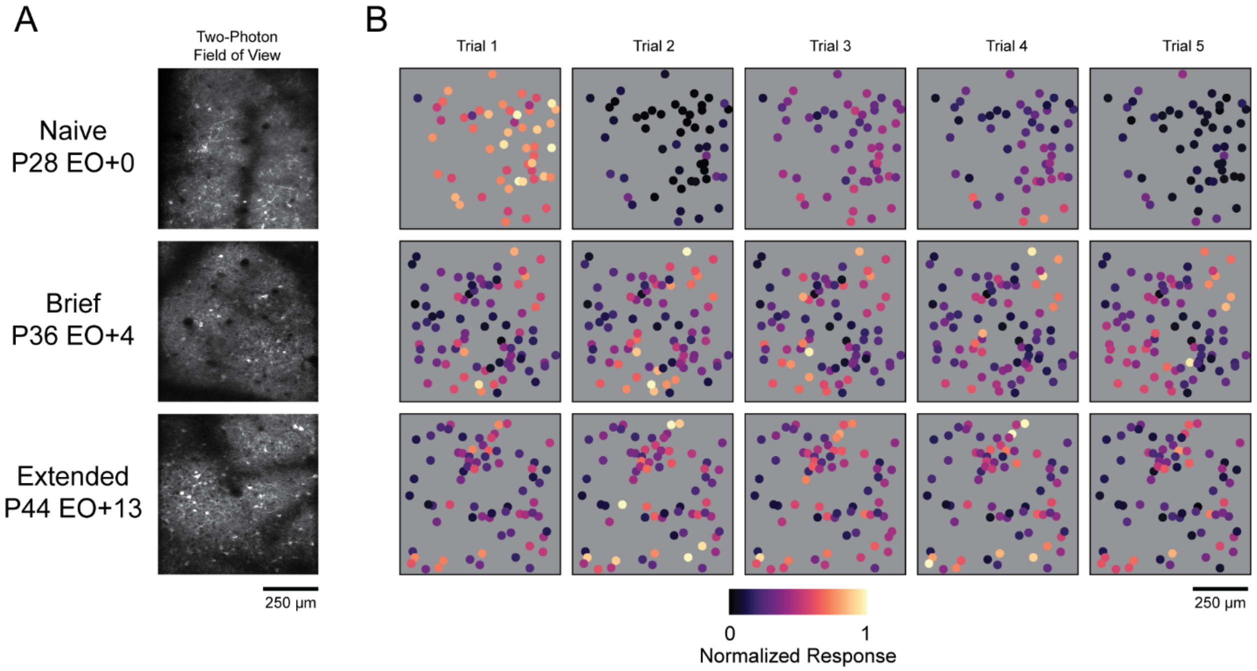


**Supplementary Figure 2 Single trial responses become more reliable with visual experience.** (A) Example two-photon fields of view for a Naive, Brief, and Extended experience animal. (B) Examples of normalized population responses for five trials of a single stimulus presentation for the corresponding animals shown in (A).

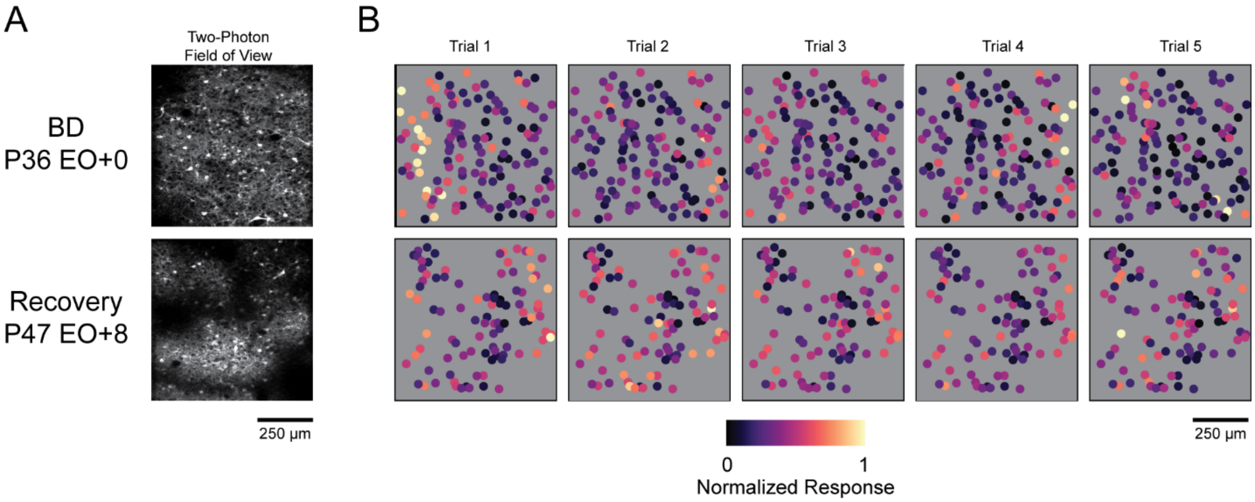


**Supplementary Figure 3 Delayed visual experience is sufficient to develop reliable responses.** (A) Example two-photon fields of view for a Binocular Deprivation and Recovery animal. (B) Examples of normalized population responses for five trials of a single stimulus presentation for the corresponding animals shown in (A).
